# Evaluating ChatGPT as an Agent for Providing Genetic Education

**DOI:** 10.1101/2023.10.25.564074

**Authors:** Nephi Walton, Sara Gracefo, Nykole Sutherland, Beth A. Kozel, Christopher J. Danford, Scott P. McGrath

## Abstract

Genetic disorders are complex and can greatly impact an individual’s health and well-being. In this study, we assess the ability of ChatGPT, a language model developed by OpenAI, to answer questions related to three specific genetic disorders: BRCA1, MLH1, and HFE. ChatGPT has shown it can supply articulate answers to a wide spectrum of questions. However, its ability to answer questions related to genetic disorders has yet to be evaluated. The aim of this study is to perform both quantitative and qualitative assessments of ChatGPT’s performance in this area. The ability of ChatGPT to provide accurate and useful information to patients was assessed by genetic experts. Here we show that ChatGPT answered 64.7% of the 68 genetic questions asked and was able to respond coherently to complex questions related to the three genes/conditions. Our results reveal that ChatGPT can provide valuable information to individuals seeking information about genetic disorders, however, it still has some limitations and inaccuracies, particularly in understanding human inheritance patterns. The results of this study have implications for both genomics and medicine and can inform future developments in this area. AI platforms, like ChatGPT, have significant potential in the field of genomics. As these technologies become integrated into consumer-facing products, appropriate oversight is required to ensure accurate and safe delivery of medical information. With such oversight and training specifically for genetic information, these platforms could have the potential to augment some clinical interactions.

## Introduction

ChatGPT is a new Large Language Model (LLM) artificial intelligence chatbot platform developed by OpenAI. ^1^ The GPT in ChatGPT stands for “Generative Pre-trained Transformer”, a type of language model that uses neural network-based transformer architecture to generate text. ^2^ The transformer architecture was introduced in a paper by Vaswani et al. in 2017^3^ and has been highly successful in natural language processing tasks. LLMs, like ChatGPT, use machine learning to generate text and are trained on a large dataset of written text, such as books, articles, and online content. They learn how to predict what word comes next in a sentence. When given a prompt or a starting point, ChatGPT uses this knowledge to generate text that is similar to the text it was trained on. Essentially, it uses statistical patterns in language to make educated guesses about what words are most likely to come next in a sentence, allowing it to generate human-like responses. LLMs have achieved state-of-the-art results on a number of natural language processing tasks and are widely used in various applications such as language translation, language generation, and text summarization. ^4,5^

The introduction of ChatGPT has received considerable media and public attention and is changing the way that scientists and clinicians think about artificial intelligence. It has demonstrated a very sophisticated ability to carry on conversations, understand human language, and answer complex questions. This technology could hold promise in its application to medical genetics and genetic counseling. Past studies have demonstrated the utility of chatbots to help answer genetic counseling questions for patients. These chatbot platforms are largely based on scripted interactions with decision trees guiding patient questions and subsequent chatbot responses. ^6-9^ There are several reasons for this, one of which is ensuring that patients receive a concrete set of predetermined information that has been not only vetted for accuracy but also specifically designed with this intent.

The introduction of ChatGPT raises the possibility that deep learning models may hold value in the future of genomics. Regardless of their current validity in the field of genomics, it is likely that consumers will be using these advanced LLMs to answer genetics questions irrespective of their adoption in formal medical applications as this platform is already seeing widespread use and is becoming part of a widely used internet search platform. ^10^ Other platforms will likely follow suit using similar technologies. Given the known shortages of genetic counselors and clinical geneticists^11-13^, patients may use these tools as an alternative to waiting for an appointment. Based on these realities, genetics professionals must pay attention to and understand these technologies to both harness their power and to understand and potentially influence how they are deployed in the public domain.

The scope of chatbots working with genetic information is rather limited, with some existing manuscripts focusing on conversational agents in the genetics and genomics space, ^6,8^ but the uniqueness of ChatGPT as a generative chatbot which uses a reinforcement learning component^2^ makes it distinct from these cases.

In this study we perform quantitative and qualitative assessments of ChatGPT’s ability to answer questions about genetic disorders related to three different genes/conditions:

*BRCA1*-related cancer syndrome, *MLH1*-related cancer syndrome, and *HFE*-related hemochromatosis.

## Methods

**IRB:** After reviewing with an Intermountain Healthcare IRB member, this study was classified as IRB exempt as it did not involve any human subjects and all queries were hypothetical in nature.

### Questionnaire design

Traditional evaluation methods for chatbots have focused on items like usability, effectiveness, efficiency user satisfaction, and esthetics. ^14,15^ However, there are a lack of validated tools used to review clinical responses. An additional challenge is that ChatGPT is a newer generative AI chatbot compared to the more traditional task-based (frequently asked questions (FAQs) or booking agent bots), rule-based (the earliest chatbots), or retrieval-based chatbots. ^16^ The difficulty in building and training generative chatbots significantly limits their appearance in published studies. After reviewing the available tools, the BOT Usability Scale (BUS-15) ^17^ was selected as the principal tool to analyze ChatGPT responses given that it contains variables designed to evaluate the satisfaction with conversational agents.

The BUS-15 is a 15-question validated survey meant to assess and compare the satisfaction of users after interactions with conversational chatbots. ^18^ It covers a range of themes related to accessibility, quality of chatbot functions, quality of conversation and information provided, perceived privacy and security, and response time. Of the 15, we chose the 7 questions that most aligned with the study goals (to review how the chatbot responds to medical prompts, particularly questions related to genetics) to be completed by domain experts (two genetic counselors and two clinical geneticists). The seven themes incorporated from BUS-15 are as follows:

- **Recognition and facilitation of users’ goal and intent**: Chatbot seems able to recognize the user’s intent and guide the user to its goals.
- **Relevance of information**: The chatbot provides relevant and appropriate information/answer to people at each stage to make them closer to their goal.
- **Maxim of quantity**: The chatbot responds in an informative way without adding too much information.
- **Resilience to failure**: Chatbot seems able to find ways to respond appropriately even when it encounters situations or arguments it is not equipped to handle.
- **Understandability and politeness**: The chatbot seems able to understand input and convey correct statements and answers without ambiguity and with acceptable manners.
- **Perceived conversational credibility**: The chatbot responds in a credible and informative way without adding too much information.
- **Meet the neurodiverse needs**: Chatbot seems able to meet needs and be used by users independently form their health conditions, well-being, age, etc.

We designed and added five additional questions to the initial seven selected from BUS-15 to evaluate the clinical value of the responses. The additional questions focused on

1. Completeness of the answer (Likert 1-7) 2) whether the response was objectively correct (Y/N), 3) if the reviewer designated the answer incorrect, what did ChatGPT get wrong (free text), 4) whether the response was lacking any key elements (free text), and 5) if anything was particularly notable in the response (free text). The BUS-15 questions used a 7-point Likert scale (Strongly Disagree to Strongly Agree). Each reviewer received a link to the survey and did not discuss the questions or answers until after survey submission.

### Use of the ChatGPT tool to create patient to AI-GC interactions

In the first stage, one clinical geneticist and two genetic counselors were tasked with conversing with ChatGPT regarding a scenario surrounding the identification of a variant in one of 3 genes (BRCA1, MLH1, or HFE). Each specialist was assigned a single gene to ask ChatGPT about. The investigators were asked to provide at least 20 freeform queries of their own design, emulating patient questions in a genetic counseling session about their respective gene. In addition to “basic” questions that the conversants believed could be easily minable from the internet (inheritance pattern, risk number), care was taken by each specialist to include questions with less objective or value driven responses (e.g., should I have children?) and those that might require interpretation of a wider scope of data (e.g., whether a human could inherit a variant from a pet?). Questions containing unintentional errors (i.e., misspellings, addressed as statements rather than questions) were retained in the study and the chatbot response to error was evaluated.

### Review of the AI-GC responses by clinical reviewers

In the second stage, resultant chatbot responses were graded using the survey instrument. For each of the 68 responses, the review started with one BUS-15 question (assessing quality) followed by the 4 clinical value questions. The remaining six BUS-15 questions were placed at the end of the survey after the evaluators had an opportunity to complete the 68 ChatGPT queries. Each response was independently rated by two genetic counselors and two clinical geneticists. Three of the reviewers had also asked questions to ChatGPT. These reviewers evaluated all the responses, including those they had asked and those asked by other participants.

The domain experts were provided both the question and the AI generated response for the clinical review.

## Statistics

The survey data were collected via Google forms. Responses were coded and transferred into R Studio (Version 2022.12.0+353) and evaluated with R (version 4.2.2) for summary statistics and conducting a Kruskal-Wallis test.

## Results

The interactions with ChatGPT 3.5 (Dec. 2022 model) resulted in 3 chat transcripts, with 21, 20, and 27 questions respectively. The full date and time stamped transcripts can be accessed in **supplemental table 1**.

Full survey results can be seen in **supplementary table 2**. The cumulative scores for the survey responses are shown below (Table 1, Likert scale: 1=strongly disagree to 7=strongly agree):

**Table 1.** Bot Usability Assessment.

| Assessment Question | Median Score |
| --- | --- |
| The chatbot provides a complete answer incorporating the information you would incorporate into the answer if asked the question. | 4.55 (Neutral/Somewhat agree);<br>*n* = 260, Range: 1-7 |
| The chatbot seems able to recognize the user's intent and guide the user to its goals. | 5.75 (Agree)<br>*n* = 4, Range: 5-6 |
| The chatbot provides relevant and appropriate information/answer to people at each stage to make them closer to their goal. | 5.75 (Agree)<br>*n* = 4, Range: 5-6 |
| The chatbot responds in an informative way without adding too much information. | 4.75 (Somewhat agree)<br>*n* = 4, Range: 4-7 |
| The chatbot seems able to find ways to respond appropriately even when it encounters situations or arguments it is not equipped to handle | 4 (Neutral)<br>*n* = 4, Range: 2-5 |
| The chatbot seems able to understand input and convey correct statements and answers without ambiguity and with acceptable manners. | 5.5 (Somewhat agree/Agree)<br>*n* = 4, Range: 5-6 |
| The chatbot responds in a credible and informative way without adding too much information. | 5.5 (Somewhat agree/Agree)<br>*n* = 4, Range: 4-7 |
| The chatbot seems able to meet needs and be used by users independently from their health conditions, well-being, age, etc. | 5.5 (Somewhat agree/Agree)<br>*n* = 3, Range: 4-5 |

To assess the reliability of the ChatGPT provided responses, we asked the reviewers whether they identified errors in each of the 68 answers (Table 2).

**Table 2.** Identification of errors in ChatGPT response*.

| Errors reported by reviewers | Count (n=68) | % of total |
| --- | --- | --- |
| None found errors (0% Yes) | 27 | 39.71% |
| Very few found errors (25% Yes) | 10 | 14.7% |
| A few found errors (33% Yes) | 7 | 10.29% |
| Split (50% Yes/50% No) | 9 | 13.23% |
| Many found errors (66% Yes) | 2 | 2.94% |
| Majority found errors (75% Yes) | 1 | 1.47% |
| All found errors (100% Yes) | 12 | 17.64% |

\*Not every question received an answer from all four respondents.

Thirty-nine percent of the 68 question responses (27 responses) were found to be error-free by all four reviewers, whereas at least 3 of the reviewers categorized 44 of the 68 questions (65%) as error-free. The remaining 24 questions (35%) were marked as containing errors by at least two respondents, with the raters unanimously scoring 12 questions as error-containing.

A Kruskal-Wallis test shown in **Figure 1** was performed on how reviewers scored the ChatGPT answers to assess if reviewers provided statistically different answers.

**Figure 1.**
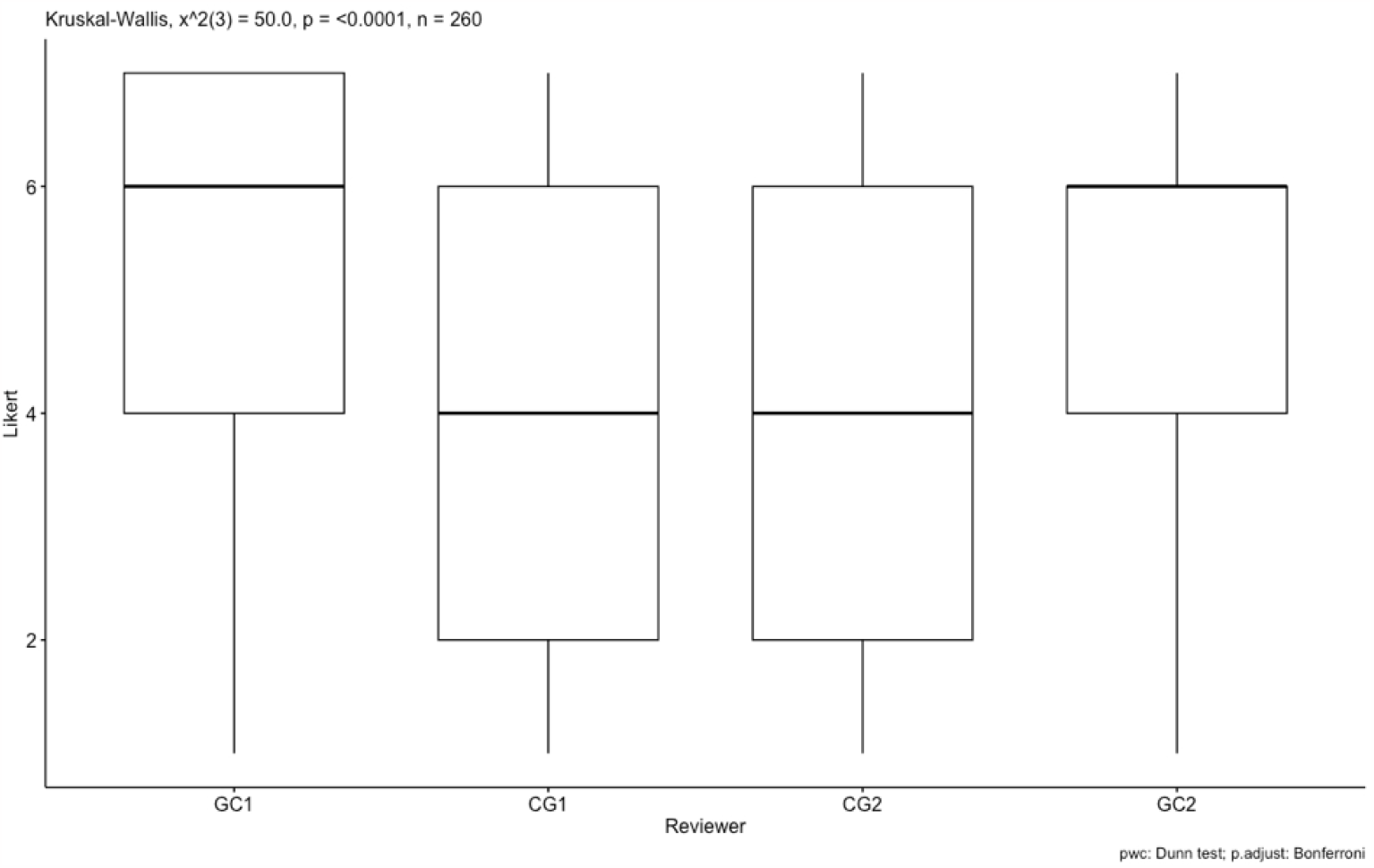
Kruskal-Wallis test assessment of reviewer responses.

A large effect size was detected (eta2[H] = 0.18). Reviewers 2 and 3 (clinical geneticists, CG1, CG2) were found to be statistically significantly different from Reviewers 1 and 4 (genetic counselors, GC1, GC2). Overall, the genetic counselors tended to rate the bot responses more favorably (1 = Strongly Disagree, 7 = Strongly Agree) than the clinical geneticists.

## Discussion

AI and ChatGPT have shown great potential in many areas^19,20^ and have also generated concern about their misuse or abuse. ^21,22^ Our study aimed to evaluate ChatGPT’s ability to effectively and accurately deliver genetic information. The results showed that ChatGPT can coherently respond to complex questions about three common genetic conditions, in some cases approaching the level of what would be delivered in a clinical setting. However, there were several weaknesses and inaccuracies identified, highlighting the need for caution before using these technologies in the medical and genetic counseling field.

The ability of ChatGPT to understand complex multipart questions and provide multi step answers was noteworthy, as shown with Question 17 (see **Figure 2)**. This is particularly significant since this chatbot was not trained specifically for the purpose of delivering genomic information. The chatbot demonstrated an extraordinary ability to carry on a conversation about the same gene or condition in the same session even if the gene or condition was not included in subsequent questions (Question 26). It gave a very accurate assessment of CRISPR in relation to genetic conditions and provided a human-like response (Questions 15 and 66). In several cases the chatbot provided additional information beyond a rote answer to the question, suggesting a deeper understanding of the goal of the question (Question 16). One response, in particular, demonstrated good insight into the potential motivations and hesitations people might have in sharing their personal health information with loved ones, and also acknowledges how sharing with family members could be a source of emotional support (Question 50). Responses like this demonstrated non-directive answers that assist the patient in weighing pros and cons without telling them what to do, which is an important concept in genetic counseling. ^23^ The chatbot was able to interpret intent, even with spelling errors (Question 24). The chatbot also responded appropriately to a statement that was not written as a question, recognizing user intent (Question 66).

One noted strength of the chatbot was its ability to converse similarly to a human providing well-written, pertinent, understandable, and convincing responses to complicated questions. Two of the reviewers noted that this capability caused them to second guess their own knowledge when they thought the chatbot was wrong, prompting verification efforts by the reviewers to prove the chatbot was indeed providing incorrect information. However, this feature may be dangerous when the chatbot provides incorrect information to patients or providers without specific content-level expertise. Within the 35% of study questions that were incorrectly answered, there were fallacies identified that could be harmful to patients (Table 2). It was remarkable that the chatbot often seemed to acknowledge its own limitations by instructing the conversant to discuss their case with medical professional to help them evaluate the recommendations it was providing (∼65% of the ChatGPT responses referenced “your doctor” or “genetic counselor”). However, this may not be a sufficient warning if the information appears valid to a lay person.

The chatbot demonstrated several weaknesses. When we looked at the 12 questions unanimously scored “incorrect” by the reviewers, no predominant themes emerged. There were some significant errors found, like the bot suggesting the question-asker should not to have children (question 37), being incorrect about dominant versus recessive inheritance (questions 25, 34, 40), appearing to misunderstand the query (question 59) or not providing a cogent answering to the query (question 32, 68). ChatGPT had some difficulty with inheritance patterns and, in particular, was unable to differentiate between autosomal recessive and autosomal dominant conditions (Question 25). This means that it is not yet capable of performing accurate risk assessment even in simple terms. Risk assessment is often complicated by family structures, such as half-relatives, and becomes even more complex with more distant relatives. It was also unable to correctly deliver recommendations based on the age of the affected person (Question 59). Of note, both deficiencies can be addressed by adding some rules to the chatbot that are attached to routinely updated knowledge bases. It was surprising that the Chatbot was able to infer that genetic conditions were not passed from humans to dogs (Question 18), however even this response had error in incorrectly stating that dogs do not have the *BRCA1* gene. Dogs do in fact have this gene and are also predisposed to mammary cancer when there are pathogenic mutations in it. ^24^

There was concern by several evaluators regarding eugenics, given the chatbot was recommending that individuals not have children based on their carriage of a recessive genetic mutation without providing other options or considerations (Question 37). Interestingly, the response to questions around childbearing differed by gene. Another response included a more correct and thorough evaluation of options (Question 5), leaving more latitude or decision making to the individual commencing in the chat. The inconsistency in the recommendations indeed makes it more human-like than providing rote responses, but also has the potential to introduce pertinent error. Asking a clinical provider the same question twice would likely result in inconsequential rephrasing, but receiving different responses to a repeated question from a chatbot could be confusing to users seeking medical answers. There were some responses that were just verifiably wrong (Question 32), and it is unclear what data may have been used to drive this response. Because the source of the data isn’t clear, it may be difficult to train a bot to only review only “trusted” sources, especially in a field that is rapidly growing. This is an example of the danger in freeform AI without rules constraining responses. It’s crucial to note that web scraping for training datasets carries the risk of including medical misinformation that is readily accessible, while authentic scientific knowledge may be inaccessible behind paywall-protected academic journals. This could lead to a bias in the training data used by ChatGPT.

Our study found that genetic counselors and geneticists had differing opinions of chatbot responses, with genetic counselors rating them more favorably. While geneticists and genetic counselors have distinct roles in patient care, potentially leading to different perspectives, the meaning of this discrepancy is unclear. Our sample size is small and additional work will be needed to understand how different members of the genetics community will respond to this new technology.

Our results show that there is still significant error in responses and that challenges remain in deploying this technology. Some of these challenges are identified in the limitations section below. However, this particular chatbot was not designed to answer questions about medical conditions, and a platform using the same technology that is trained specifically for this task and given rules around genes/disease (autosomal dominant versus recessive inheritance etc.) would likely perform markedly better. As we documented in our study, ChatGPT can provide what appears to be well constructed answers, even when they are wrong (questions 25, 34, 40). AI generated answers from these search engines may prove even more compelling than the hyperlink-based results from the past, increasing the possibility of people using these results to determine medical care.

From our findings, two aspects emerge that are very important. One is AI’s role in the future of delivering genomic care. The other is the expectation that AI will be incorporated in future search engines and consumers will be using this technology to ask complicated questions, whether society deems it ready or not. In the short term, perhaps the latter situation is more concerning. As chatbots are integrated into consumer search engines, there is concern about the accuracy of information delivered. More research is needed to determine how best to handle this issue as chatbots become more convincing in their ability to interpret and answer questions. The fact that they “seem” convincing with interaction makes them more dangerous if they are delivering incorrect information. Delivery of erroneous information may cause many negative sequalae, including leading consumers to make misinformed decisions in their healthcare, contribute to mistrust in medical care, and even bypass the medical care that they need. ^25,26^

Overall, we believe that the technology behind ChatGPT has shown some momentous results and is extremely powerful with the ability to change the world much like the internet has. This is generating a lot of excitement. With “great power comes great responsibility”, ^27^ and those who deploy AI systems that have the capability to deliver medical information, regardless of intended use, must put in place protections to ensure they are not leading patients down harmful paths. We must not forget that the knowledge and experience that this model is built upon relies largely on human-generated information. While AI is touted to have the capability to start generating such information on its own, doing so without significant human oversight brings up many concerns. AI models for genomics must be updated constantly or at least Iteratively, to ensure they are up to date. This makes regulation difficult as the learning models are not static and will produce different results depending on the date the question is asked.

As we assess ChatGPT for a potential role in delivering education about genes and diseases to the lay population, it is important to note that genetic counseling is much more than an information exchange. Generative AI has potential as a tool to assist with information provision, but it remains limited in its ability to provide counseling as it is unable to tailor responses to individuals’ unique situations and circumstances. AI, at least in the near term, is not capable of supplanting human interaction when it comes to fully communicating what it means to have a genetic condition. More importantly, AI is unlikely to ever provide the compassion and emotional support that is crucial for those going through this experience.

## Limitations

Our study had several limitations. First, there are a lack of a validated instruments for evaluating chatbot responses to clinical questions. Future potential work in this area could involve the development of such a tool, in addition to continuing to test and monitoring LLM’s ability to assist in medical information dissemination. Second, we tested ChatGPT on relatively common genetic conditions about which there is a large amount of minable information present on the internet. Further studies should probe the ability of this algorithm to adequately perform in rarer or newly discovered conditions, given the rapidly evolving field.

## Conclusion

AI is showing significant growth and has the potential to supplant humans in certain applications. This is as concerning as it is hopeful. There is tremendous potential for such AI platforms in genomics that are specifically trained to deliver genomic information in any language. As more generalized technologies, such as ChatGPT, are incorporated into consumer-facing products, there must be some oversight to ensure that they are not delivering misinformation regarding medical care. This becomes difficult in a world where there is so much uncertainty, and opinions differ even amongst specialists. Regulatory guiderails should be established that encompass different viewpoints and opinions in care while ensuring that these platforms do not deliver directly harmful information. With proper oversight and training specifically for genetic information, platforms like ChatGPT could have the potential to augment some human interactions in the field of genomics.

## Data Availability

The full dataset with both questions asked to ChatGPT, its responses, and survey responses are attached in the appendix (supplementary tables 1 & 2).

## Acknowledgments

The authors would like to thank Melanie McGrath PhD LAT ATC for assistance in the preparation of the manuscript.

## Funding Statement

BAK’s effort was supported by the Intramural Research Program of the NIH and NHLBI. All other costs associated with the research at other institutions were borne by the authors.

## Author Contributions

NW and SPM designed the study. NW, SG, and NS developed the queries submitted to ChatGPT. BAK, NW, SG, and NS rated the ChatGPT responses. SPM performed the data analysis. All authors contributed to the manuscript.

## Ethics Declaration

All questions created for querying ChatGPT were hypothetical and were crafted by the domain experts based off their experience working in the medical genetics field. No human subjects were used in this study.

## Conflict of Interest

The authors declare that they have no conflicts of interest regarding the publication of this paper.

## Notes

### Competing Interest Statement

The authors have declared no competing interest.

